# A Single-Dose Bundibugyo Virus Vaccine Protects Macaques Within 3 Days

**DOI:** 10.64898/2026.06.14.732188

**Authors:** Kyle L. O’Donnell, Jil A. Haase, Corey W. Henderson, B. Ryder Gathright, Paige Fletcher, Joseph F. Rhoderick, Chad S. Clancy, Shilpi Jain, César Albariño, Brian J. Smith, Andrea Marzi

## Abstract

Bundibugyo virus (BDBV), a member of the orthoebolaviruses in the *Filoviridae* family, causes severe hemorrhagic disease with high case-fatality rates. Currently, there are no medical countermeasures approved for human use hampering the response to the large ongoing outbreak in the Democratic Republic of the Congo and Uganda. While vesicular stomatitis virus (VSV)–based vaccines have demonstrated fast-acting prophylactic single-dose efficacy against multiple filoviruses, it has yet to be defined for VSV-BDBV. Here, we evaluated the rapid protection by a single-dose vaccination of VSV–BDBV in nonhuman primates (NHPs). Vaccination elicited rapid innate and early adaptive immune responses and conferred complete protection from clinical disease against BDBV challenge within 3 days. Vaccinated NHPs exhibited minimal clinical signs, limited systemic inflammation, and no infectious virus was isolated from the blood at any time. Protection correlated with neutralizing antibodies and Fc effector functionality of the humoral immune response. These findings establish VSV–BDBV as a fast-acting vaccine candidate suitable for outbreak response and highlight immune mechanisms underlying rapid protection.

## Introduction

Bundibugyo virus (BDBV), a member of the orthoebolaviruses, causes severe hemorrhagic disease referred to as Ebola disease (EBOD) with high case-fatality rates; no approved medical countermeasures (MCMs) are currently available ^1^. BDBV emerged in 2007 in western Uganda and caused a major EBOD outbreak with a ∼40% case fatality rate ^2^. Currently, the largest BDBV outbreak is ongoing in the Democratic Republic of the Congo (DRC) and Uganda. To date there have been 400 confirmed cases and 66 deaths, including healthcare workers. This has led the World Health Organization to declare the outbreak a global health emergency ^3,5^. Due to the unpredictable outbreak capacity of filoviruses, there is an unmet need for MCMs which are rapidly effective.

The nonhuman primate (NHP) model remains the gold standard for assessing filovirus MCMs. The clinical manifestation of EBOD in NHPs accurately reflects what is observed in humans including high viremia, hypercytokinemia, coagulopathy, which may progress to disseminated intravascular coagulopathy, multiorgan failure, and death^6^. Unlike other models of EBOD with Ebola virus (EBOV) or Sudan virus (SUDV) which are uniformly lethal, BDBV does not cause uniform lethality in cynomologus macaques ^6–8^. This is reflective of the decreased pathogenic nature in humans in comparison to the other orthoebolaviruses ^9^.

Recombinant vesicular stomatitis virus (VSV) vaccines have been successfully deployed against EBOV, demonstrating excellent efficacy in ring-vaccination strategies and rapid onset of protection ^10–13^. However, cross-protection between filovirus species is limited, necessitating virus-specific vaccine constructs ^14–16^.

Vaccines conferring rapid onset of protection are particularly critical for filovirus outbreak containment where transmission occurs through close contact with infected bodily fluids and where healthcare-associated exposures are common. While VSV-based vaccines can elicit protective immunity within 7–10 days in NHPs against EBOV, SUDV, and MARV the immunologic correlates and feasibility of achieving this rapid protection against BDBV remain unknown ^17–21^. Previous studies showed that a recombinant VSV vaccine expressing the full-length BDBV glycoprotein (GP; VSV-BDBV) was uniformly protective when administered 29 days prior to challenge, and partially protective when administered 20-30 mins post-exposure ^7,8^. Here, we sought to determine if vaccination 3 or 7 days before challenge would result in uniform protection. We demonstrate that a single dose of the VSV–BDBV vaccine induces complete protection within 3 days. We characterized innate and adaptive immune responses underlying this rapid protection and add to the growing body of evidence that VSV-based vaccines are fast-acting MCMs highlighting the use of a VSV-BDBV vaccine in BDBV outbreaks.

## Results

### VSV–BDBV vaccination protects NHPs within 3 days

To determine a shortened time from vaccination to challenge resulting in uniform protection from disease caused by BDBV infection, cynomolgus macaques (n=5 per group) received a single intramuscular (IM) dose of VSV–BDBV (1×10^7^ PFU) either 7 or 3 days prior to IM challenge with 10,000 median tissue culture infectious dose (TCID_50_) of BDBV. A control group received the VSV-Marburg virus (VSV-MARV) vaccine at 1×10^7^ PFU by the IM route 7 days prior to BDBV challenge. Control vaccination resulted in 60% lethality (Fig. 1A); 4/5 NHPs of the control group demonstrated with clinical scores indicating moderate to severe disease (Fig.1 B). Serum chemistry analysis revealed that the control NHPs developed increased levels of aspartate aminotransferase (AST) and blood urea nitrogen suggestive of a multiorgan dysfunctional disease phenotype (Fig.1 C,D). Viral load quantification showed that the control NHPs developed high titer viremia, specifically the NHPs that succumbed to disease, while the VSV-BDBV-vaccinated NHPs displayed no or low viral RNA levels (Fig. 1 E); no infectious virus was recovered at any time point from the blood of all vaccinated NHPs (Fig. 1F). Control NHPs also demonstrated irregular patterns of viral shedding assessed through the collection of swabs measuring viral RNA and infectious viral titers (Supplemental Figure 1); BDBV RNA was absent in all swab samples obtained from the vaccinated NHPs (Supplemental Figure 1). Tissue viral loads were assessed to determine the extent of systemic viral spread into peripheral organs. The control NHPs that succumbed to infection had high levels of viral RNA in the tissues collected at necropsy (8-9 days post-challenge, DPC) including multiple lymph nodes, liver, spleen, adrenal gland, and gastroduodenal junction; all typical target tissues of systemic filovirus spread (Fig. 1G). The control NHPs that survived infection displayed reduced viral replication in the tissues at the time of euthanasia (42 DPC) similar to the vaccinated groups (Fig. 1H). Residual viral RNA was detected in the vaccinated groups at the time of euthanasia (42 DPC); levels were absent or reduced compared to the control NHPs that succumbed to acute disease (Fig. 1H).

**Figure 1:**
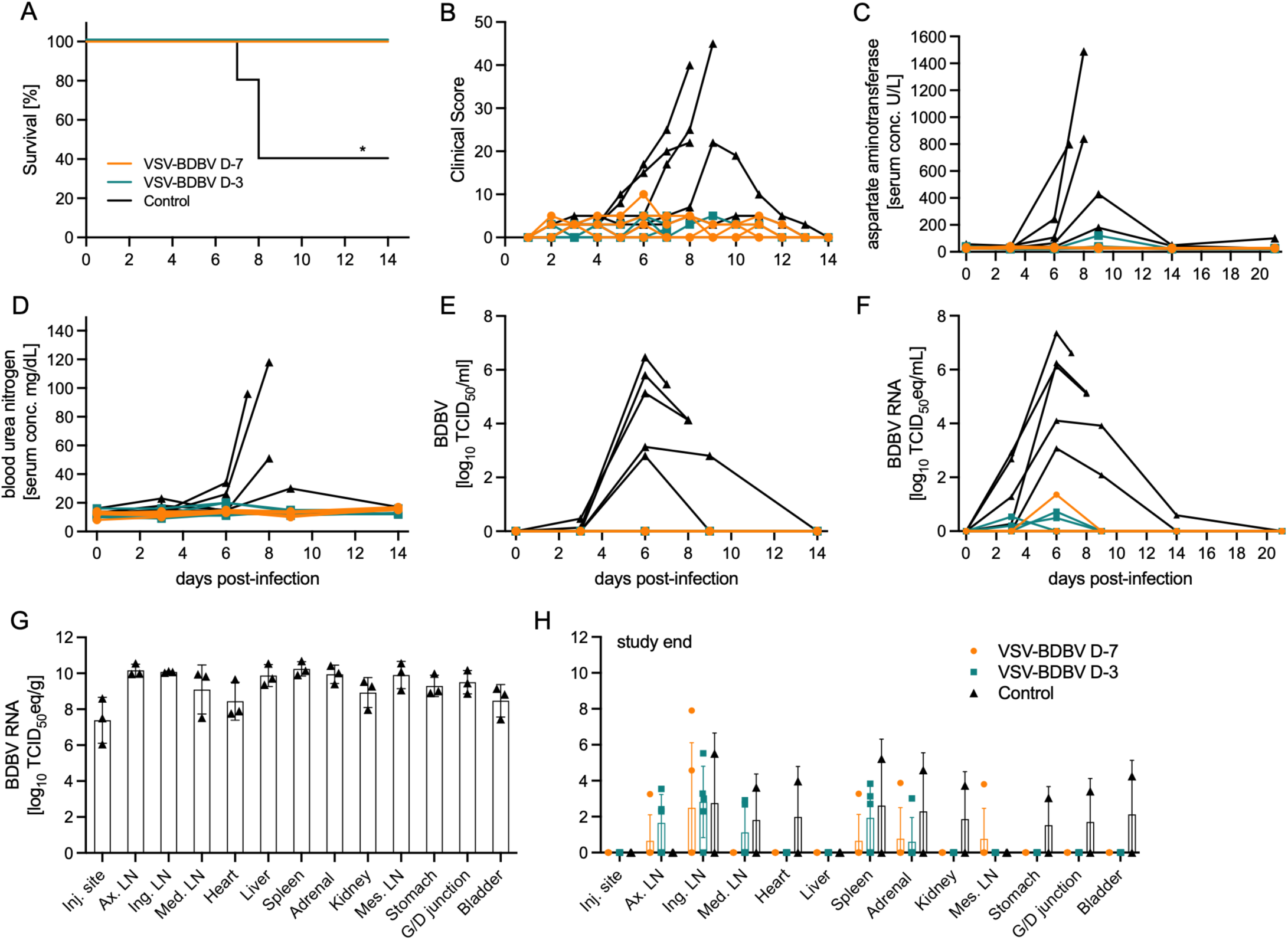
Clinical data of vaccinated NHPs challenged with BDBV. Groups of 5 NHPs were vaccinated at the indicted time and infected with BDBV on day 0. (A) Survival and (B) clinical score of infected NHPs. Changes in serum chemistry through the acute disease phase of the study included changes in (C) aspartate aminotransferase and (D) blood urea nitrogen. Levels of (E) infectious BDBV or (F) BDBV RNA in the blood. BDBV RNA in the tissue of control NHPs that succumbed to disease (G); BDBV RNA samples in survivors at day 42 (H). Data sets were analyzed using (A) Log-rank Mantel-Cox test; statistically significant differences are indicated in comparison to the control group as *p*<0.05 (*). TCID_50_, median tissue culture infectious dose; eq, equivalent; Inj. injection; Ax. LN, axillary lymph node; Ing. LN, inguinal lymph node; Med. LN, mediastinal lymph node; Mes. LN, mesenteric lymph nodes; G/D junction, gastroduodenal junction.

### VSV-BDBV vaccination prevents EBOD pathology

Histopathologic evaluation of the samples obtained from control NHPs showed classic signs of EBOD in both the liver and spleen including moderate hepatocellular necrosis and inflammation centered along portal triads and vasculitis of the central vein (Fig. 2 A,D). In comparison, VSV-BDBV-vaccinated NHPs had no observable histopathologic changes observed at 42 DPC (Fig. 2B,C,E,F). Similarly, the spleen of control NHPs showed mild to moderate neutrophilic and necrotizing splenitis with accumulation of degenerate and non-degenerate neutrophils along the periphery of lymphoid follicles whereas no lesions were observed in VSV-BDBV vaccinated NHPs (Fig. 2D-F). Further systemic disease was observed in control NHPs including arcuate arteritis with or without renal pelvis hemorrhage in the kidney (Fig. 2G). Multiorgan pathology in the control NHPs included necrotizing and hemorrhagic adrenalitis, neutrophilic and necrotizing interstitial pneumonia with or without alveolar fibrin, pulmonary edema and alveolar exudates, necrotizing lymphadenitis, germinal center necrosis with accumulation of histiocytes in subcapsular sinuses in all examined peripheral lymph nodes (Supplemental Figure 2). Interestingly, in evaluated complete sections of the epididymis of control NHPs (n=4), the head of the epididymis exhibited single cell epithelial necrosis, interstitial epididymis, vasculitis and/or interstitial hemorrhage (Supplemental Figure 2). Similarly, gastric hemorrhage was observed in the lamina propria of 3/5 control NHPs (Supplemental Figure 2).

**Figure 2:**
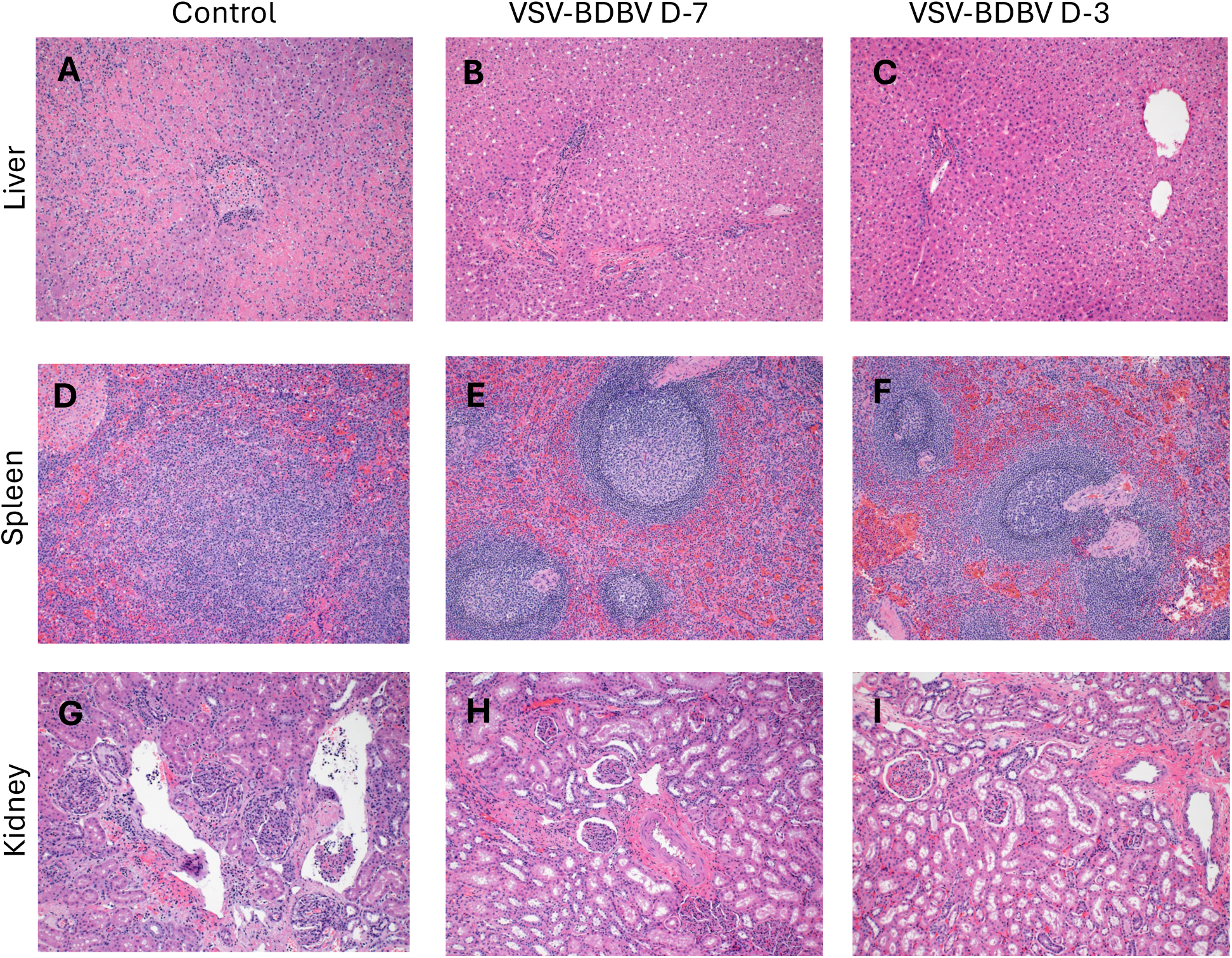
Histopathologic findings in NHPs after BDBV Challenge. (A) Severe centrilobular hepatocellular necrosis with leukocyte infiltration and central vein vasculitis. (B,C) Liver with no significant histopathologic changes. (D) Moderate splenic follicular necrosis and lymphoid disorganization. (E,F) Normal splenic lymphoid population and architecture. (G) Interstitial nephritis, hemorrhage and arcuate artery vasculitis. (H, I) Normal kidney architecture. Hematoxylin and eosin staining imaged at 200x.

### Humoral immunity drives rapid protection

The primary immune-mediated protective mechanism stimulated by VSV-based vaccines is the rapid induction of the humoral response ^20,22,23^. Therefore, we assessed the antigen-specific humoral response and functionality in detail. BDBV GP-specific IgG was detected within one week in the NHPs receiving the vaccine 7 days prior to challenge (Fig. 3A). Peak IgG titers were measured on 9 DPC and remained elevated for the duration of the study (Fig. 3A). Interestingly by 14 DPC the surviving control NHPs had similar BDBV GP-specific IgG titers compared to the vaccinated NHPs. The functional capacity of the humoral response was also assessed and the NHPs receiving the vaccine 7 days prior to challenge developed a more robust functional antibody profile with higher amounts of neutralizing capacity, phagocytic capabilities, and NK cell activation markers compared to the control NHPs at 0 DPC (Fig. 3B-H). Correlation analysis on 6 DPC, which corresponds to a peak disease time point, revealed that overall IgG titers, neutralization titers, cellular phagocytosis, and NK cell activation markers correlate with improved disease outcomes (Supplemental Figure 3, Supplemental Table 1). This evidence further reinforces the importance of a multifunctional humoral response to be generated for protection from EBOD.

**Figure 3.**
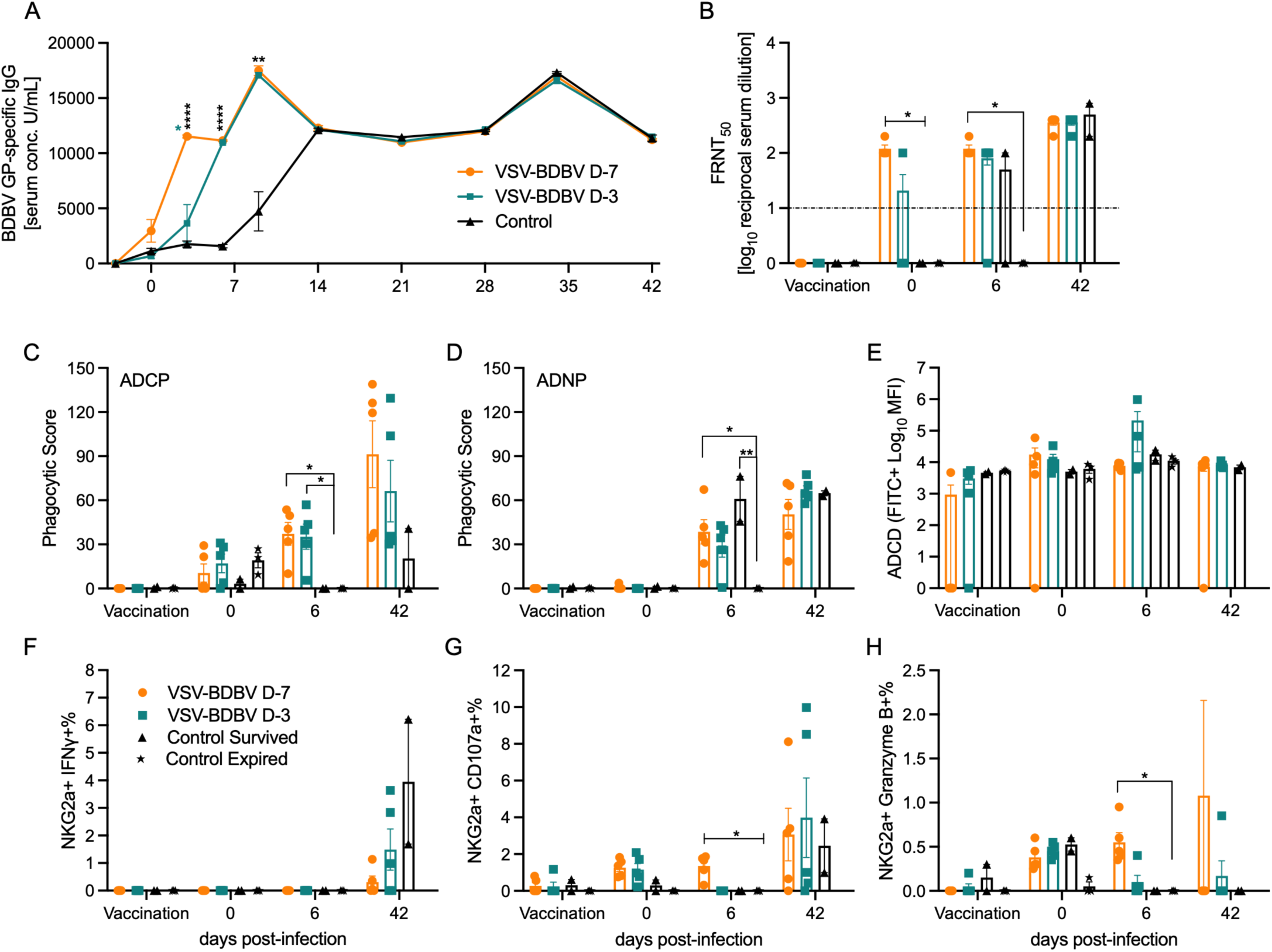
Humoral immune responses after vaccination and BDBV challenge. BDBV GP-specific (A) serum IgG titers were determined over time; control NHP terminal samples during acute disease were grouped with day 9. (B) Serum neutralization presented as FRNT_50_ of GFP-positive cells at the time of vaccination, challenge (day 0), day 6, and study end (day 42). Dotted line indicates assay limit of detection. (C) Antibody-dependent cellular phagocytosis, ADCP; (D) antibody-dependent neutrophil phagocytosis, ADNP, and (E) antibody-dependent complement deposition, ADCD, activities were determined in serum. Antibody-dependent NK cell activation, ADNKA, markers (F) IFNγ, (G) CD107a, and (H) granzyme B are presented as percent positive of the total NK cell population. Mean (SE; A, B, C, D, F, G, and H) and geometric mean (SD; E) are depicted. Data sets were analyzed using Kruskal-Wallis test with Dunn’s multiple comparisons, statistically significant differences are indicated as p < 0.0001 (****), p < 0.01 (**), or p < 0.05 (*).

### T cell and cytokine responses aid in BDBV control

While the primary protective immune response of VSV-based vaccines is the humoral response, the cellular response does contribute to the protective profile ^17,20^. Using flow cytometric analysis, we determined that the activated central and effector memory CD4 T cell compartments were both expanded in the VSV-BDBV-vaccinated groups (Supplemental Figure 4). This was also the case for central memory CD8 T cells demonstrating that the cellular response being activated is likely contributing to the refinement of the humoral response as well as acting as a cytotoxic CD8 T cell response controlling viral spread (Supplemental Figure 4). These changes, however, did not reach a level of significance but highlight immunologic trends that may be contributing to protection. Consistent with a benign clinical disease course, vaccinated NHPs exhibited minimal cytokine dysregulation following challenge. Pro-inflammatory cytokines characteristic of severe EBOD, including IL-6, TNF-α, IFN-α, IFN-β, and IFN-γ, were markedly reduced compared to the control NHPs (Fig. 4A-E). Further examples of a dysregulated immune response are the significant increases in IL-10 and IP-10 associated with the NHPs that succumbed to disease (Fig. 4F,G).

**Figure 4.**
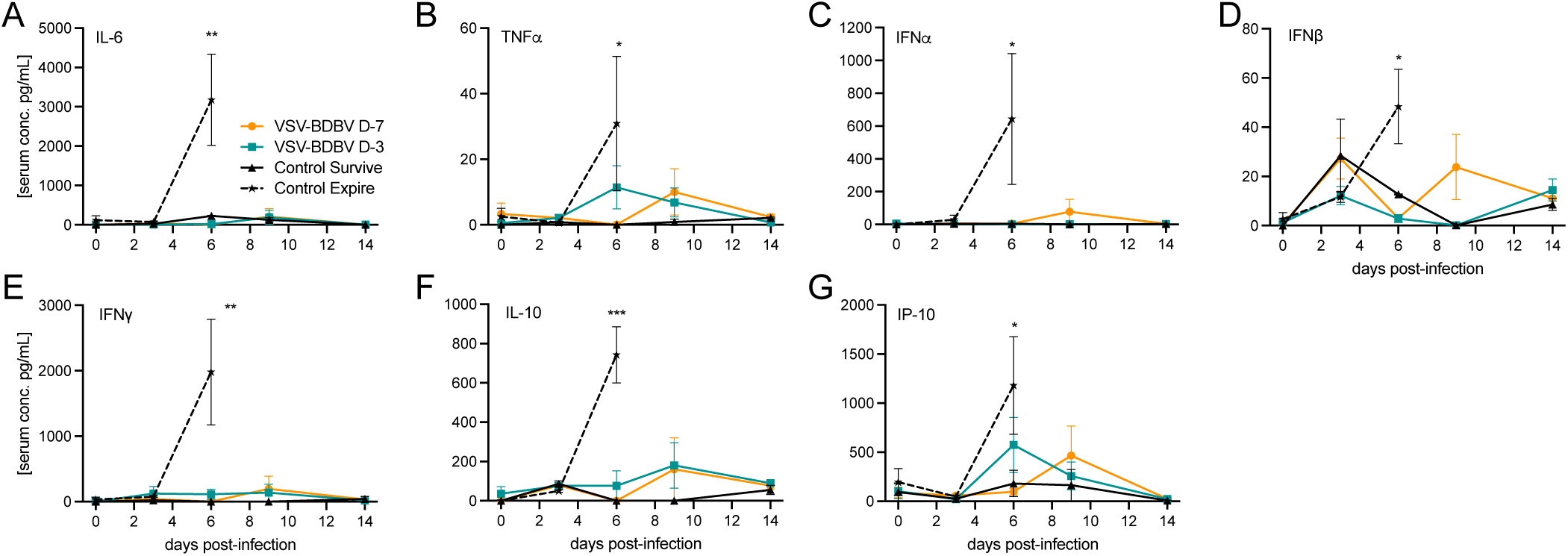
Cytokine responses in vaccinated NHPs after BDBV challenge. Kinetics of IL-6, TNFα, IFNα, IFNβ, IFNγ, IL-10, and IP-10 in serum samples collected after BDBV challenge. Mean and standard error of the mean are depicted. Data sets were analyzed using Kruskal-Wallis test with Dunn’s multiple comparisons; statistically significant differences are indicated as p < 0.001 (***), p < 0.01 (**), or p < 0.05 (*).

## Discussion

Here, we demonstrate that a single dose of VSV–BDBV confers complete protection when administered as early as three days prior to a partially lethal BDBV challenge by IM exposure. The challenge virus used in this study has 98.8% sequence identity with the contemporary strain in the DRC and Uganda further validating our data and their applicability to the current outbreak ^24^. The rapid onset of protection observed here is comparable to that reported for the licensed VSV–EBOV vaccine (Ervebo by Merck; rVSV-ZEBOV) as well as the VSV-SUDV and VSV-MARV vaccines currently in clinical development ^18–20^. Given the unpredictable nature of filovirus outbreaks, MARV, BDBV and SUDV retain a significant outbreak potential with a complete lack of approved MCMs to rapidly deploy in the event of emergence. An additional public health concern is that in the event of an outbreak, such as the one currently ongoing in the DRC and Uganda caused by BDBV, public health officials may be pressured into implementing all strategies at their disposal. One such strategy may be the use of approved vaccines for other orthoebolaviruses which have historically resulted in poor outcomes in preclinical models ^19,25^. The filovirus field urgently needs to expand the very limited preclinical data available in this context to ensure proper guidance for MCM deployment in future outbreaks.

The demonstration of rapid onset of protection for species-specific VSV-based vaccines against the human-pathogenic filoviruses indicates that this may be an intrinsic property of the VSV platform dominated early on by the robust innate immune response. The protective efficacy of VSV-based filovirus vaccines altogether, however, is species-specific, driven primarily by the antibody responses directed against the viral GP antigens. The GPs of different orthoebolaviruses exhibit significant antigenic variation, therefore the licensed EBOV vaccine has limittaions if a virus from another species causes an outbreak. Our group has previously demonstrated that the VSV-EBOV showed no protection against heterologous SUDV challenge in NHPs ^19^. While the timing between vaccination and heterologous challenge seem to impact the disease course, there is limited cross-reactivity and no cross-protective efficacy between EBOV and SUDV^15,19^. In the context of BDBV challenge, VSV-EBOV has been assessed in numerous formulations such as a single vaccination, mixture with VSV-SUDV, and as a prime-boost strategy with VSV-SUDV as the prime vaccine and VSV-EBOV as the boost vaccine. The single immunization approach with VSV-EBOV provided 75% protective efficacy (3/4 NHPs) ^26^. The VSV mixture did not provide any protective benefits compared to the control group, while the prime-boost strategy resulted in 100% survival similar as the VSV-BDBV vaccine ^7^. Due to the inherent nature of the BDBV challenge model not being uniformly lethal, additional NHP studies are needed to accurately confirm the cross-protective phenotype. Taking this further, staggered vaccinations against filoviruses using even different vaccine platfoermds may result in an unkown cross-protective potential that warrants investigation in preclinical models.

In this study, rapid protection appears to be associated with the induction of early antiviral innate immune responses coupled with the accelerated development of functional GP-specific antibodies. Control NHPs received the same dose of the VSV-MARV vaccine shown to elicit robust innate immune repsonses ^21^ but did not survive the BDBV challenge highliting that indeed the antigen-specific immune response is critical for protection. In contrast, NHPs that succumbed to disease exhibited evidence of a dysregulated innate immune response following challenge, characterized by significantly elevated levels of type I and type II interferons, TNFα, IL-6, and IL-10. This dysregulated pro- and anti-inflammatory cytokine response is characteristic of human and NHPs suffering from severe EBOD ^29,30^. Future studies are needed to define the cellular sources of cytokine production and to determine how their activation states contribute to disease progression.

The humoral response is strongly associated with protection at early time points, particularly at 6 DPC, when both vaccine groups exhibit significantly higher IgG titers compared to control NHPs. While antibody binding titers are traditionally the main indicator of immunity and neutralization is the conventional functional readout of antibody activity for many viruses—and has been shown to contribute to protection even at low levels—our data highlights the importance of assessing additional antibody functions. Consistent with this, neutralizing titers remained modest at 6 DPC, whereas Fc-mediated antiviral functions—previously implicated in EBOV vaccine-mediated protection—were robust and likely contributed to viral clearance. Among these, antibody-dependent cellular phagocytosis (ADCP) was elevated in both vaccine groups as early as 6 DPC compared to control NHPs. Additionally, antibody-dependent NK cell activation (ADNKA) differed significantly between animals vaccinated 7 days prior to challenge and those that succumbed to disease, as measured by increased expression of CD107a and granzyme B. Together, these markers are indicative of NK cell-mediated cytotoxic activity against infected cells. The delayed response observed between vaccination groups may reflect differences in the time required for affinity maturation of antigen-specific IgG and the acquisition of effective FcγRIII engagement, although this remains to be directly tested. To determine if humoral functionality was a true correlate of protection in this model, we implemented Spearman’s correlation analysis. The correlation analysis showed that ADCP, ADNKA, and neutralization titer were all strong correlates of protection regarding clinical score and body temperature. An antibody functionality profile with the combination of neutralization, ADCP, and ADNKA is ideal for a filovirus vaccine platform as it eliminates the risk of antibody-dependent enhancement through the complement activation pathway ^31^. It has also been previously shown that EBOD survivors have had high levels of neutralizing and phagocytic antibodies ^32,33^ indicating that the rapid and robust antibody response stimulated upon vaccination has a favorable functionality profile for continued protection.

Implications of the cellular immune response in VSV-based vaccinations have previously been described ^34^. Primarily a CD4 T-cell response bolsters the humoral response through Th2 type mechanisms. However, it has previously been established that orthoebolaviruses use VP24 and VP35 to inhibit type I IFN signaling which has a downstream effect on T-cell receptor signal transduction ^35^ which has previously been shown to dampen the vaccine-mediated antiviral T-cell response. Upon investigation into memory subsets of T-cells we determined that central memory CD4+ T-cells are more activated and expanding in the VSV-BDBV vaccine groups compared to the control NHPs that survived the challenge. The trend is also extended to the activated central memory CD8+ T-cells which may play a role in direct antiviral activity. This is indicative of a difference in the vaccine response compared to challenge virus response. Allowing the rapid generation of central memory T-cells to differentiate to effector memory subsets induces additional antiviral capacity. In the future additional characterization will need to be conducted on the T cell compartment to elucidate the antiviral functional capacity of these cells in detail.

A limitation of this study is that the cellular phenotyping of the early innate immune cells was not conducted. While this was not the focus of this study, cellular phenotyping or single cell RNA sequencing would provide interesting information about the early immune response and provide additional insights into vaccine-mediated protective mechanisms and correlates of protection. It has previously been demonstrated from a genomic perspective that a reduction in myeloid-derived suppressor cells is correlated with survival in this model ^36^. While we achieved significant differences in the survival between control and vaccinated NHP groups, n=5 is the minimum group size and a limitation of this study. Larger group sizes would have enabled a much broader analysis of the protective immune responses and could guide future studies towards the identification of a correlate of protection. Additionally, there is currently no assay for BDBV GP-specific IgM analysis available to us, similarly only the BDBV GP-specific responses were assessed when other VSV proteins may be playing a role in protection and should be assessed in future studies. Lastly, a control group receiving the VSV-MARV vaccine 3 days prior to challenge would have allowed us to decipher intrinsic details

These results have immediate implications for outbreak response. A fast-acting and specific BDBV vaccine would be suitable for ring vaccination, frontline healthcare worker protection, and emergency deployment during outbreaks such as the one currently ongoing in the DRC and Uganda. Future work will focus on the durability of the immunity elicited by the VSV-BDBV vaccine and protection will be evaluated in mucosal exposure models. The ability to fully protect within 3 days—despite BDBV’s antigenic divergence from EBOV—highlights the versatility of the VSV platform and provides a foundation for fast-acting multivalent or pan-filovirus vaccination strategies.

## Materials and Methods

### Ethics statement

All work involving infectious BDBV was performed following standard operating procedures (SOPs) approved by the Rocky Mountain Laboratories (RML) Institutional Biosafety Committee (IBC) in the maximum containment laboratory at the RML, Division of Intramural Research, National Institute of Allergy and Infectious Diseases, National Institutes of Health. Procedures were conducted by trained personnel under the supervision of veterinary staff on anesthetized animals. All efforts were made to ameliorate animal welfare and minimize animal suffering per the Weatherall report on the use of nonhuman primates in research (https://royalsociety.org/policy/publications/2006/weatherall-report/). Animal work was performed in strict accordance with the recommendations described in the Guide for the Care and Use of Laboratory Animals of the National Institute of Health, the Office of Animal Welfare, and the United States Department of Agriculture and was approved by the RML Animal Care and Use Committee (ACUC). Animals were housed in adjoining individual primate cages that enabled social interactions, under controlled conditions of humidity, temperature, and light (12-h light:12-h dark cycles). Food and water were available *ad libitum*. Animals were monitored and fed commercial monkey chow, treats, and fruit at least twice a day by trained personnel. Environmental enrichment consisted of commercial toys, music, and video. Endpoint criteria based on clinical score parameters as specified and approved by the RML ACUC were used to determine when animals were humanely euthanized.

### Animal study design

Fifteen male cynomolgus macaques (*Macaca fascicularis*) 6-11 years of age and 5.5–7.9 kg in weight were used for this study and randomly assigned to vaccination groups. Group sizes of n = 5 were determined to be required by power analysis using log-rank test in graph pad/prism under the assumption that 40% of the control animals may survive the infection. Two groups of cynomolgus macaques were vaccinated with a single IM injection of 1 × 10^7^ PFU VSV-BDBV and challenged 7 or 3 days later. Control NHPs were IM-vaccinated with 1 × 10^7^ PFU VSV-MARV and challenged 7 days later. An IM injection of 10,000 TCID_50_ of BDBV (confirmed by back-titration) as previously described ^13^ served as lethal challenge for all NHPs. Clinical exams including a blood draw on anesthetized NHPs were conducted on −7, −5, −3, −1, (pre-challenge exams depended on group) 0, 1, 3, 6, 9, 14, 21, 28, 35 and 42 DPC. The animals were observed daily for clinical signs of disease according to a RML ACUC-approved scoring sheet and humanely euthanized when they reached endpoint criteria. The study ended 42 DPC when all surviving animals were euthanized.

### Cells and viruses

Vero E6 cells (*Mycoplasma* negative, RRID: CVCL_0059) were grown at 37 °C and 5% CO_2_ in Dulbecco’s modified Eagle’s medium (DMEM) (Sigma-Aldrich, St. Louis, MO) containing 10% fetal bovine serum (FBS) (Wisent Inc., St. Bruno, Canada), 2 mM L-glutamine, 50 U/mL penicillin, and 50 mg/mL streptomycin (all supplements from Thermo Fisher Scientific, Waltham, MA). THP-1 (*Mycoplasma* negative, RRID:CVCL_0006) were grown at 37 °C and 5% CO_2_ in Roswell Park Memorial Institute medium (RPMI) (Sigma-Aldrich, St. Louis, MO) containing 10% FBS (Wisent Inc., St. Bruno, Canada), 2 mM L-glutamine, 50 U/mL penicillin, and 50 mg/mL streptomycin (all supplements from Thermo Fisher Scientific). HL-60 (*Mycoplasma* negative, RRID:CVCL_0002) were grown at 37 °C and 5% CO_2_ in improved minimum essentail medium (IMEM) (Sigma-Aldrich, St. Louis, MO) containing 20% fetal bovine serum (FBS) (Wisent Inc., St. Bruno, Canada), 2 mM L-glutamine, 50 U/mL penicillin, and 50 mg/mL streptomycin (all supplements from Thermo Fisher Scientific, Waltham, MA). VSV vaccines and BDBV were propagated in Vero E6 cells using DMEM supplemented with 2% FBS, L-glutamine, and penicillin/streptomycin. VSV-BDBV was constructed in-house as previously described ^37^ and used for IM vaccination. BDBV strain 200706291 (GenBank accession No. MK028856.1), was propagated and titered on Vero E6 cells, and stored in liquid nitrogen until challenge. rBDBV-ZsG was constructed as described previously ^38^. Briefly, viral RNA from BDBV isolate EboBund-122 2012 (GenBank accession KC545395.1) was used as a template to be amplified by RT-PCR. The overlapping fragments were used to assemble a full-length clone in the T7 transcription vector. The ZsGreen open reading frame and porcine teschovirus 2A self-cleaving peptide were cloned at the amino terminus of the nucleoprotein. The virus was recovered following transfection of Huh-7 cells seeded in 12-well plates with 0.5 μg pBDBV-ZsG, 0.5 μg pC-L, 0.25 μg pC-NP, 0.05 μg pC-VP35, 0.05 μg pC-VP30, and 0.5 μg of codon-optimized pC-T7. Supernatants were harvested 3 days after transfection and blind-passaged twice in Huh-7 cells. Viral stocks were sequenced by NGS on the MiniSeq platform (Illumina, San Diego, CA, USA). All viruses were confirmed by sequencing.

### Hematology and serum chemistry

The total white blood cell, neutrophil, lymphocyte, and platelet counts were determined from EDTA blood with the IDEXX ProCyte DX analyzer (IDEXX Laboratories, Westbrook, ME). Serum biochemistry including aspartate aminotransferase (AST), alkaline phosphatase (ALP), alanine aminotransferase (ALT), glucose, blood urea nitrogen, creatinine, and total bilirubin was analyzed on a Vetscan 2 using Preventive care profile disks (Abaxis, Union City, CA).

### Histopathology

Tissues were collected at the time of euthanasia when animals reached endpoint criteria or at study end (42 DPC). Tissues were fixed in 10% Neutral Buffered Formalin (2 changes), for a minimum of 7 days. Tissues were placed in cassettes and processed with a Sakura VIP-6 Tissue Tek, on a 12-hour automated schedule, using a graded series of ethanol, xylene, and PureAffin. Embedded tissues are sectioned at 5um and dried overnight at 42 °C prior to staining. Representative images of liver, spleen, kidney, lymph node, adrenal gland, lung, stomach and epididymis were captured on an Olympus BX51 microscope using an Olympus DP80 camera with cellSens Dimension software (Olympus, Center Valley, PA). All tissues were evaluated by a blinded, board-certified veterinary pathologist.

### BDBV RNA quantification

RNA from EDTA blood samples was extracted using the QIAamp Viral RNA Mini Kit (Qiagen, Hilden, Germany) according to manufacturer specifications. Tissues, a maximum of 30 mg each, were processed and RNA was extracted using the RNeasy Mini Kit (Qiagen) according to manufacturer specifications. One step RT-qPCR for genomic viral RNA was performed using a specific primer-probe set [<u>forward primer 5’-GAAGGGTCAGGTGCCTTATTAC-3’, reverse</u> <u>primer 5’-GTGTGGTTGTCCCAGAAACT-3’), FAM probe (5’-ACAGAGCAT/ZEN/AGCATCGAGGCAGAA-3’)]</u> to the BDBV L gene, and the QuantiFast Probe RT-PCR + ROX Vial Kit (Qiagen), in the Rotor-Gene Q (Qiagen). Five μL of each RNA extract were run alongside dilutions of BDBV standards with a known concentration of RNA copies. Concentrations were determined using the Q-Rex 1.1.04 software with the absolute quantification plugin.

### BDBV titers

Viremia was determined from EDTA whole blood samples using high-passage (>90 passages) Vero E6 cells (*Mycoplasma* negative). BDBV titers were determined from swab samples collected in 1 mL plain DMEM. Cells were seeded in 48-well plates the day before titration. On the day of titration, blood samples were thawed, and 10-fold serial dilutions were prepared in DMEM without supplements. Media was removed from cells and inoculated in triplicate with each dilution. After 1 h, DMEM supplemented with 2% FBS, penicillin/streptomycin, and L-glutamine was added, and cells were incubated at 37 °C. Cells were monitored for cytopathic effect (CPE) until 14 days post titration and a TCID_50_ was calculated for each sample using the Reed and Muench method ^39^.

### Antigen-specific humoral responses

Post–challenge NHP sera were inactivated by gamma-irradiation (4 MRad) and removed from the maximum containment laboratory according to SOPs approved by the RML IBC ^40^. The BDBV GP-specific IgG titers in serum samples were determined at 1:250 dilution using ELISA kits following manufacturer’s instructions (Alpha Diagnostics, San Antonio, TX).

### Quantification of antibody Fc effector functions

Assays for antibody effector functions were adapted from previously established protocols ^23,41^. Post–challenge NHP sera were inactivated by gamma-irradiation (4 MRad) and removed from the maximum containment laboratory according to RML SOPs approved by the RML IBC ^40^. Recombinant BDBV GPΔTM (IBT Bioservices) was tethered to Fluospheres NutrAvidin-Microspheres yellow-green or red (Thermo Fisher Scientific, Waltham, MA) using the EZ-link Micro Sulfo-NHS-LC-Biotinylation kit (Thermo Fisher Scientific).

### Antibody-dependent complement deposition (ADCD)

Serum samples were heat-inactivated at 56°C for 30 min then diluted 1:200 in DMEM and applied to the conjugated beads (20 μl/well) for one hour at 37°C. Next, guinea pig complement (Cedarlane, Burlington, Canada) was added for 30 minutes. The bead complexes were washed with PBS containing 15 mM EDTA and stained with anti-C3c-FITC (Antibodies-Online, Pottstown, PA; Cat. No. ABIN458597). Data were acquired on a FACS Symphony (BD Biosciences, Franklin Lakes, NJ) and analyzed in FlowJo v10.

### Antibody-dependent neutrophil phagocytosis (ADNP)

HL-60 cells (4×10^5^ cells/mL) were differentiated with 1.3% DMSO for 5 days 37 °C and 5% CO_2_. Biotinylated BDBV GPΔTM (IBT Bioservices) was coupled to yellow-green Neutravidin beads (Life Technologies). Serum samples were diluted 1:250 in culture medium and incubated with GP-coated beads for 2 hours at 37 °C. Beads (20 μL) were added to HL-60 cells (5×10^4^ cells/well) and incubated for 2 hours at 37 °C. Cells were then stained for CD11b (Clone G10F5; BioLegend), CD16 (Clone UCHT1; BD Biosciences), and fixed with 4% paraformaldehyde and resuspended in cell staining buffer (Biolegend). Data were acquired on a FACS Symphony (BD) and analyzed in FlowJo v10. Neutrophils were defined as SSC-A^high^ CD11b^+^, CD16+. A phagocytic score was determined using the following formula: (percentage of FITC^+^ cells)*(geometric mean fluorescent intensity (gMFI) of the FITC^+^ cells)/10,000.

### Antibody-dependent cellular phagocytosis (ADCP)

Serum samples were diluted 1:250 in DMEM and incubated with 20 μl of the conjugated beads for 2 hours at 37°C. The serum/bead mixture was then transferred to a plate of THP-1 cells (2.5×10^4^ cells/well) and incubated overnight at 37°C. Cells were fixed with 4% paraformaldehyde and resuspended in cell staining buffer (Biolegend, San Diego, CA). Data were acquired on a FACS Symphony (BD) and analyzed in FlowJo v10. A phagocytic score was determined using the following formula: (percentage of FITC^+^ cells)*(geometric mean fluorescent intensity (gMFI) of the FITC^+^ cells)/10,000.

### Antibody-dependent natural killer cell activation (ADNKA)

Recombinant BDBV GPΔTM (IBT Bioservices) was coated onto MaxiSorp 96-well plates (Thermo Fisher Scientific) at 300 ng/well at 4 °C overnight. Wells were washed with PBS and blocked with 5% BSA prior to addition of serum and incubation for 2 hours at 37 °C. Unbound antibodies were removed by washing with PBS, and NK92 cells expressing CD16 were added at 5×10^4^ cells/well in the presence of 4 μg/mL brefeldin A (Sigma-Aldrich), 5 μg/mL GolgiStop (Thermo Fisher Scientific) and anti-CD107a antibody (Clone H4A3, BD Biosciences) for 5 hours. Cells were surface stained for CD16 (clone 3G8, Pacific Blue; Biolegend Cat. No. 302032) and CD56 (clone 5.1H11; Alexa Fluor488; Biolegend Cat. No. 362518) for 20 min. Cells were fixed and permeabilized with Fix/Perm (Life Technologies) according to the manufacturer’s instructions to stain for intracellular IFNγ (Clone B27, PE; Biolegend Cat. No. 506507) and TNFα (clone Mab11, APC; Biolegend Cat. No. 502912). Data were acquired on a FACS Symphony (BD Biosciences) and analyzed in FlowJo v10.

### Neutralization

The day before this assay, Vero E6 cells were seeded into 96 well plates. Serum samples were heat-inactivated at 56 °C for 30 min and 5-fold serially diluted in DMEM. BDBV-ZsGreen was added in equal volumes at a MOI of 3, and the mixture was incubated for 1 h at 37 °C. The antibody-viral solution was then transferred to the cells and incubated for 24 h at 37 °C and 5% CO_2_. The cells were fixed with 4% PFA and resuspended in cell staining buffer (Biolegend). Sample acquisition was performed on a Cytoflex-S (Beckman Coulter, Brea, CA) and data analyzed in FlowJo v10.

### Serum cytokine quantification

Serum samples were diluted 1:4 in assay buffer for analysis using the BioLegend Legendplex NHP anti-virus response panel as per the manufacturer’s instructions (Biolegend, San Diego, CA). Sample acquisition was performed on a FACS Symphony (BD). Concentrations for IFN-γ, IL-6, TNF-α, IP-10, IFN-α, IFN-β, and IL-10 were determined for all samples. Values below the limit of detection of the assay were assigned the value of 1.

### Cellular phenotyping assays

PBMCs were isolated from whole blood samples using Histopaque®-1077 (Sigma-Aldrich) and separated according to manufacturers’ instructions. Isolated PBMCs were resuspended in FBS with 10% DMSO and frozen at −80 °C until analysis.

For T cell response analysis, cells were stimulated in duplicate with 2 μg/mL BDBV GP peptide pool, media, cell stimulation cocktail (containing PMA-Ionomycin, Biolegend), or SARS-CoV-2 nucleocapsid peptide pool together with 5 μg/mL Brefeldin A (Biolegend) for 16 h. Following, cells were surface stained with: CD11b-BV605 (BD), CD45-Alexa700 (BD) CD3-PerCP-Cy5.5 (BD), CD4-BUV496 (BD), CD8-BV570 (BD) CD95-PE-Cy5 (Biolegend), HLA-DR-BV650 (BD), FoxP3-BV421 (BD), Ki-67-FITC (BD), CD20-APC-Cy7 (BD), IgG-PE (BD), IgM-BV786 (BD) CD28-PE-CF594 (BD), NKG2a-Pe-Cy7 (Beckman Coulter) CD16-BV711 (Biolegend), CD56-BUV661 (BD), Live/Dead-aqua (Biolegend). Sample acquisition was performed on a FACS Symphony (BD) and data analyzed in FlowJo V10. The gating strategy is depicted in Supplemental Figure 5. All flow cytometry antibodies were validated by the production companies performing antibody titrations on cells expressing the respective proteins with proper control cells to ensure consistency between production lots ^42,43^.

### Statistical analysis

All statistical analysis was performed in Prism 11 (GraphPad). Statistical significance of survival was determined by log rank Mantel-Cox test. All other data were evaluated by Kruskal–Walis test with Dunn’s multiple comparisons or two-tailed Mann–Whitney test. Statistical significance was achieved at p < 0.05 and is indicated in each figure. Spearman correlation analyses between IgG endpoint titers and functionality readout and clinical symptoms were performed using the JMP® statistical analysis software. The *X*–*Y* scatterplots show 95% confidence density ellipses for normally distributed data. The table indicates Spearman correlation coefficients that were statistically significant (*p* < 0.05).

## Acknowledgements

We are grateful to all members of the Rocky Mountain Veterinary Branch, National Institute of Allergy and Infectious Diseases (NIAID), National Institutes of Health (NIH), particularly the animal caretakers, for their support of this study. This research was supported by the Intramural Research Program of the National Institutes of Health (NIH) (ZIA AI001254 to AM). The contributions of the NIH and CDC authors are considered Works of the United States Government. The findings and conclusions presented in this paper are those of the authors and do not necessarily reflect the views of the NIH, the CDC or the U.S. Department of Health and Human Services.

## Author contributions

A.M. conceived the idea, designed the study, and secured funding. K.L.O., C.W.H., C.S.C, B.J.S. and A.M. conducted the animal study. K.L.O., J.A.H., C.W.H., B.R.G., P.F., C.S.C., J.F.R. and A.M. processed samples, performed assays, and analyzed the data. S.J. and C.A. generated the BDBV-GFP virus and shared it with A.M. under an MTA. K.L.O. and A.M. wrote the manuscript with input from all authors. All authors approved the manuscript.

## Correspondence

Correspondence should be addressed to Andrea Marzi.

## Data availability

Data supporting the findings of this study are available at figshare link.

---

**Supplemental Figure 1.**
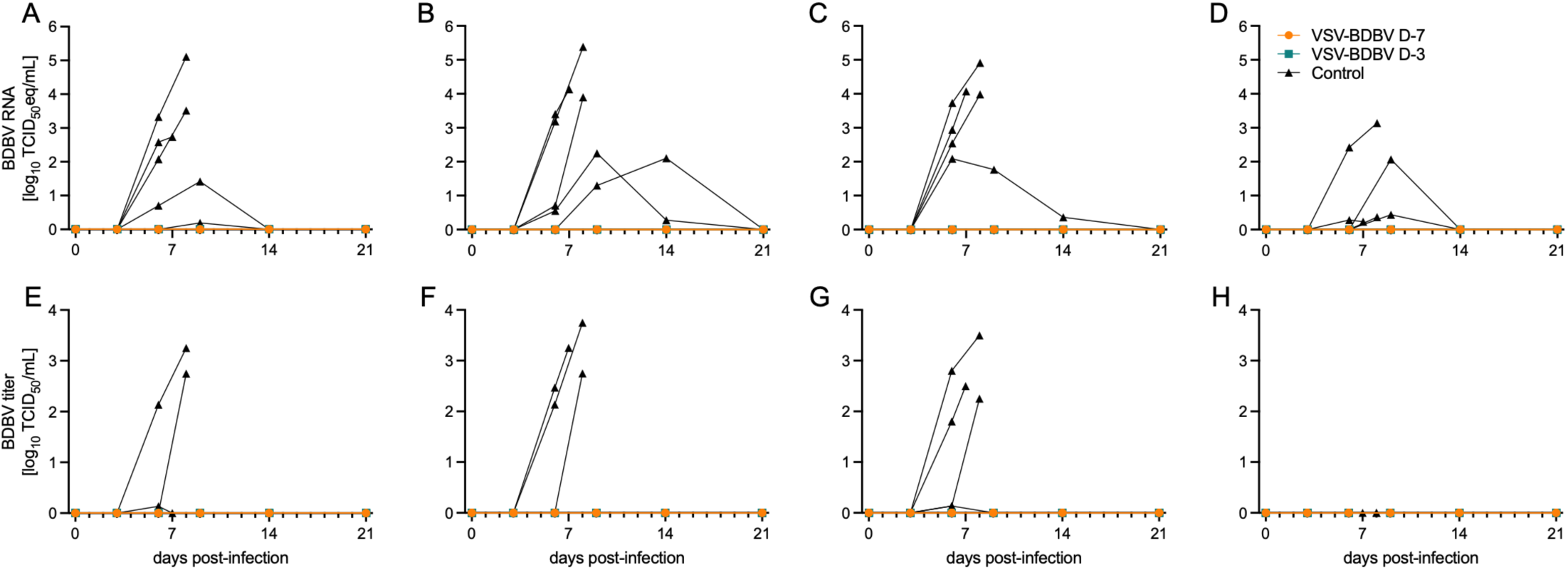
Viral shedding dynamics in BDBV-infected NHPs. BDBV RNA in the (A) oral, (B) nasal, (C) rectal, and (D) urogenital swabs. BDBV titer in the (E) oral, (F) nasal, (G) rectal, and (H) urogenital swabs. TCID_50_, median tissue culture infectious dose; eq, equivalent;

**Supplemental Figure 2.**
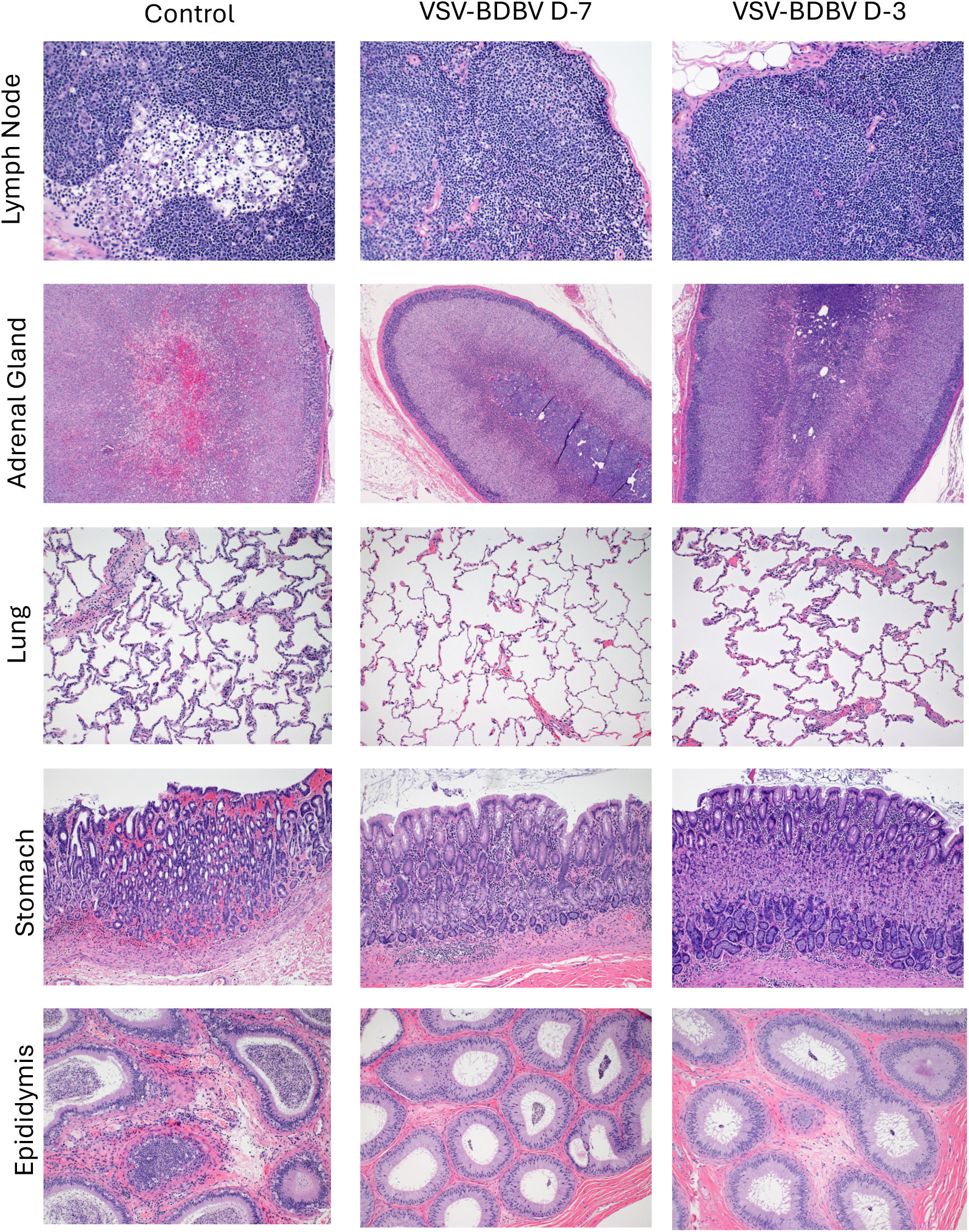
Systemic histopathlogic findings in NHPs after BDBV challenge. Lymph node, adrenal gland, lung, stomach and epididymis samples collected at the time of necropsy at acute disease (control) or at study end. Hematoxylin and eosin staining imaged at 200x.

**Supplemental Figure 3.**
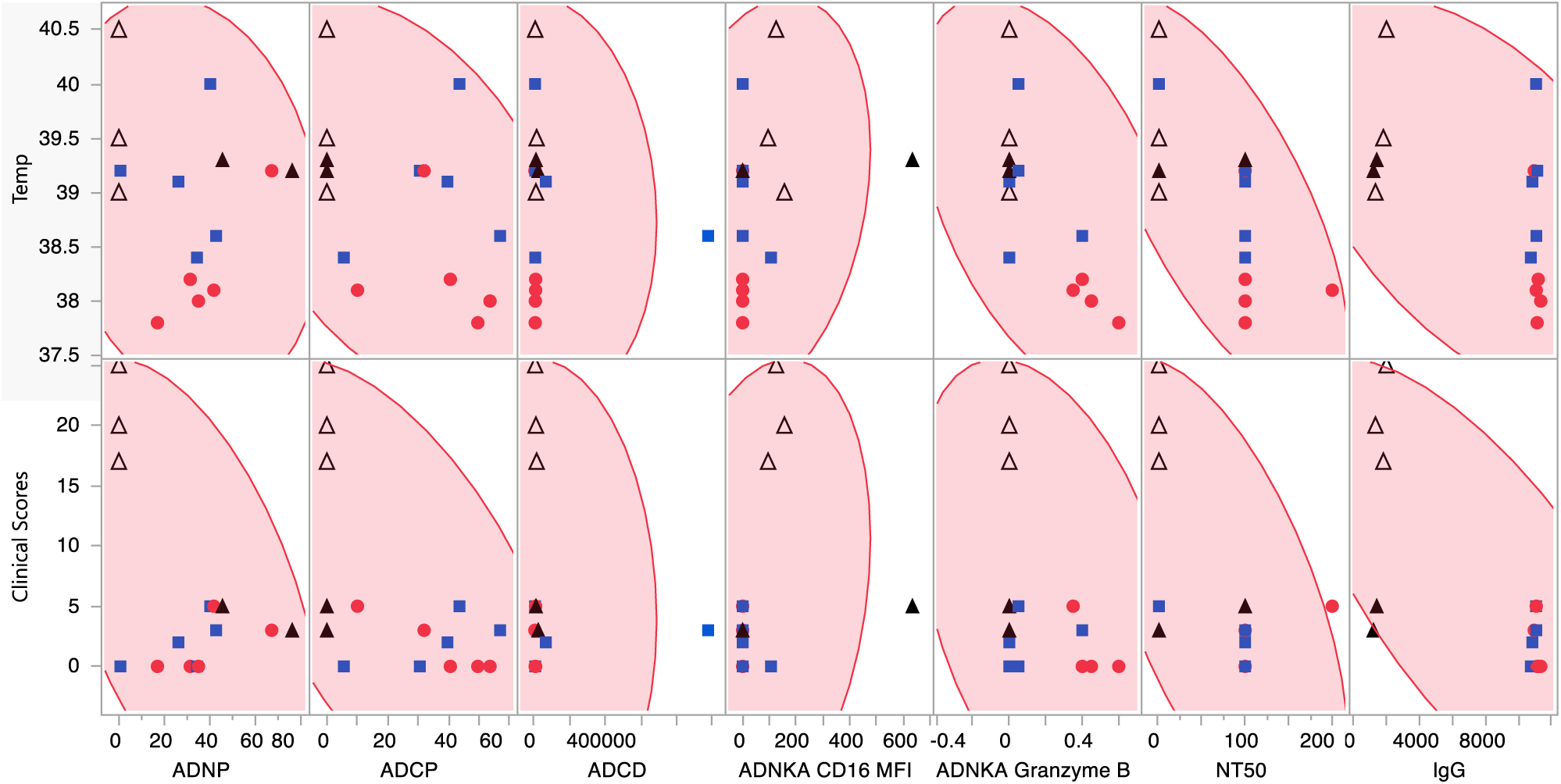
Correlation analysis of survival outcomes and antigen-specific humoral responses. Scatterplot matrix showing relationship between clinical symptoms (body temperature and clinical score) and immunogenicity readouts measured on 6 days post-challenge (n=5 per group). Data sets (provided in Supplemental Table s) were analyzed using Spearman’s test (two-tailed).

**Supplemental Figure 4.**
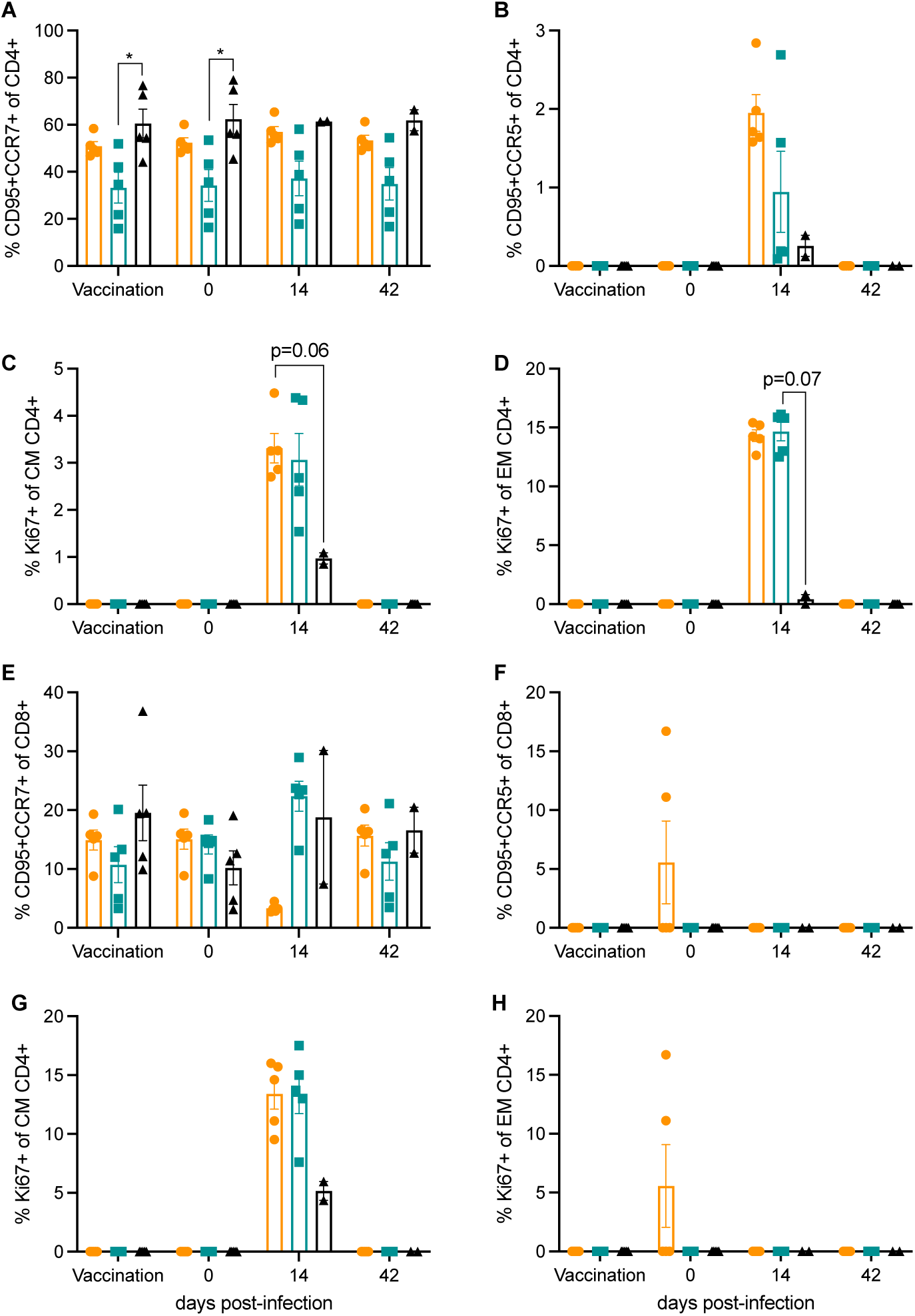
Cellular immune response after vaccination and BDBV challenge. (A-D) CD4 T cells and (E-H) CD8 T cells were characterized for memory populations and activation markers were determined over time. (A) Central memory (CM) CD4 T cells; (B) Effector memory (EM) CD4 T cells; (C-D) activation of CM and EM with Ki67 staining. (E) Central memory (CM) CD8 T cells (F); effector memory (EM) CD8 T cells; (G-H) activation of CM and EM with Ki67 staining. Mean (SE) are depicted. Data sets were analyzed using Kruskal-Wallis test with Dunn’s multiple comparisons, statistically significant differences are indicated as p < 0.05 (*).

**Supplemental Figure 5.**
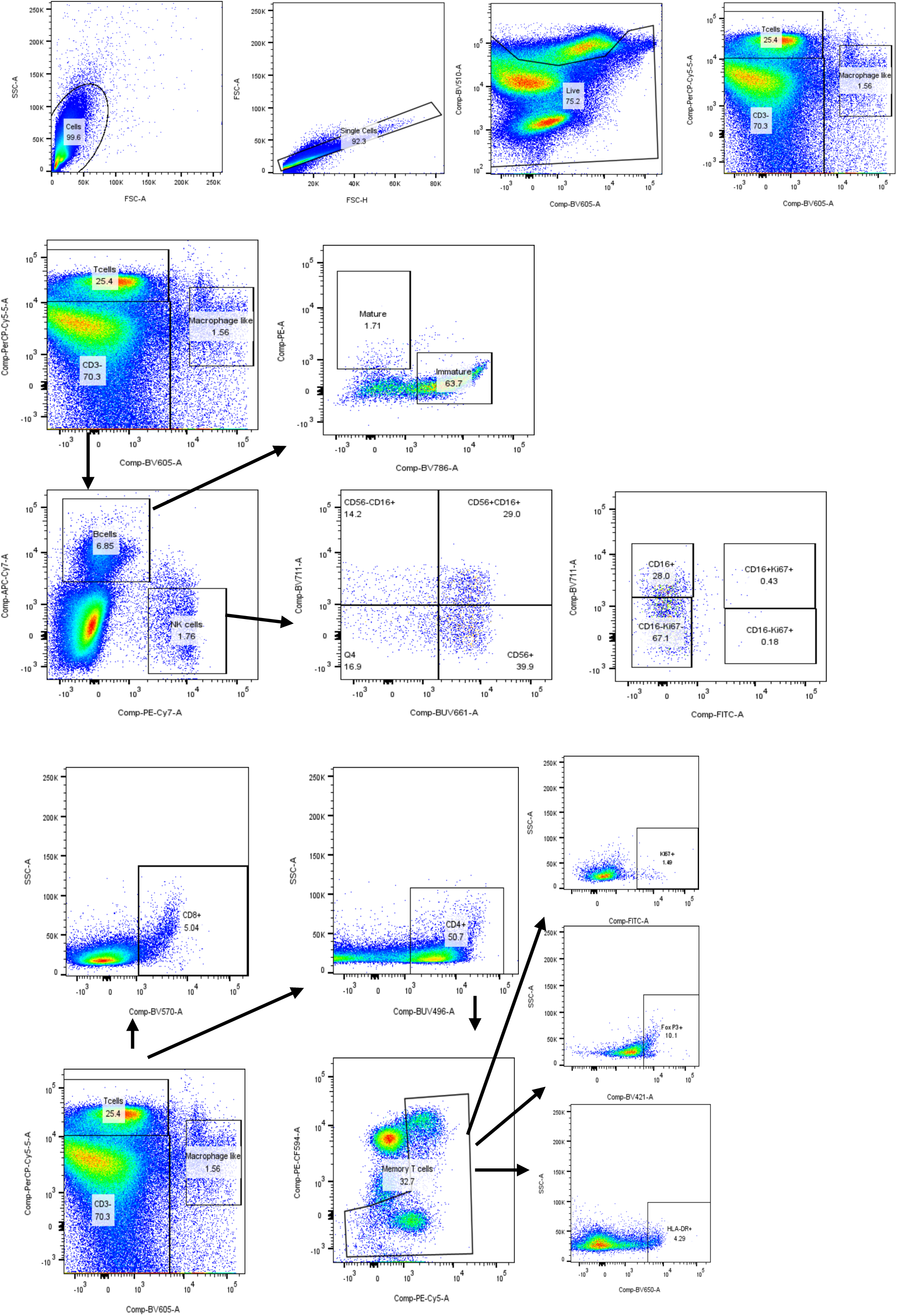
Flow cytometry gating strategies.

**Supplemental Table 1.** Spearman analysis of clinical outcomes and antigen-specific humoral responses 6 DPC. Spearman correlation coefficients (ρ) and two-tailed p-values highlighting relationships that are statistically significant (p < 0.05). No adjustments were made to the data prior to analysis.

| Variable | by Variable | Spearman $\rho$ | Prob>\|$\rho$\| |
| --- | --- | --- | --- |
| Clinical Scores | ADCD | 0.164889378 | 0.557032618 |
| Clinical Scores | ADCP | -0.59703155 | 0.01878077 |
| Clinical Scores | ADNKA CD16 MFI | 0.554088776 | 0.032092424 |
| Clinical Scores | ADNKA Granzyme B | -0.477957497 | 0.071546051 |
| Clinical Scores | ADNP | -0.14341725 | 0.610111667 |
| Clinical Scores | IgG | -0.66322172 | 0.007033743 |
| Clinical Scores | NT50 | -0.55850037 | 0.030467972 |
| Temperature | ADCD | -0.0143 | 0.9596 |
| Temperature | ADCP | -0.5293 | 0.0425 |
| Temperature | ADNKA CD16 MFI | 0.3841 | 0.1575 |
| Temperature | ADNKA Granzyme B | -0.5338 | 0.0404 |
| Temperature | ADNP | -0.0719 | 0.7989 |
| Temperature | IgG | -0.5627 | 0.029 |
| Temperature | NT50 | -0.6593 | 0.0082 |

